# TET1 Functions as an Oxidized LDL Dependent Early-Stage Inducer of Atherosclerosis by Initiating Foam Cell Formation in Macrophages

**DOI:** 10.64898/2026.02.04.703872

**Authors:** Anvitha Boosani, Jonathan A. Green

## Abstract

Atherosclerosis is a condition characterized by plaque growths in arteries, consisting of oxidized LDL (low-density lipoprotein) and localized cell cumulation. By the time of diagnosis for patients with atherosclerosis, the disease has often progressed into advanced stages. Statins are commonly prescribed; however, while these drugs can lower blood cholesterol levels, they cannot regress or stop the plaque growth. Currently, there are no treatments available to prevent the formation of new plaques. Such treatment options would require the identification of proteins that act during disease onset, initiating molecular mechanisms that promote plaque formation.

Histone deacetylases (HDACs) and Ten Eleven Translocation (TET) demethylases are two important classes of epigenetic mediators. Some isoforms of these two classes of proteins have been found to transcriptionally regulate cellular inflammation, which may favor plaque formation. These transcriptional regulators seem to function early in the molecular mechanisms that are involved in disease progression. In the present work, we identified a clear role of these epigenetic proteins in foam cell formation. Foam cells have been implicated as part of the early steps which ultimately lead to atherosclerosis. Here we showed that in the presence of OxLDL (oxidized LDL), the protein isoform TET1 has a direct role in foam cell formation, while HDAC2 adopts a more indirect role. Using specific inhibitors of TET1 and HDAC2, we showed the inter-regulated molecular mechanisms between these proteins and how they regulate foam cell formation *in vitro*.

In this study, we found that upon inhibition of TET1 in U937-derived macrophages, and subsequent foam cell formation via OxLDL treatment, a lower percentage of foam cells was observed. However, TET2 inhibition under the same treatment conditions had no effect on the inhibition of foam cell formation.

## 1 Introduction

Cardiovascular disease (ischemic heart disease) is the leading cause of death in the world. Atherosclerosis is the most prevalent type of cardiovascular disease. Atherosclerotic plaques begin as “fatty streaks,” or small deposits of OxLDL within the medial layer of the artery. These become plaques and grow when the cellular mechanisms that remove the lipid buildup fail. The response to the accumulation of OxLDL in the arterial wall initially begins with monocytes that enter the vicinity, transform into macrophages, and attempt to efflux the OxLDL. Ultimately, these macrophages become saturated and remain within the OxLDL deposit. These cells are now known as foam cells, and they recruit more immune cells to the area, thus causing more foam cells to form and accumulate. This influx of macrophages causes the plaque to grow. As these plaques grow, they begin to obstruct blood flow, leading to life-threatening complications.

Although there is evidence suggesting that vascular smooth muscle cells can also engulf OxLDL, it is primarily macrophages that attempt to efflux OxLDL and eventually transform into foam cells ^1–4^. When exposed to OxLDL, monocytes (the primary cell types that transition into macrophages) can induce the production of several proatherogenic mediators, including IL6 and TNF-α. The inflammatory responses triggered by these proatherogenic proteins contribute to increased foam cell formation. Many of these steps are mediated through intracellular proteins that initiate epigenetic mechanisms ^5^. The molecular mechanisms initiated by the buildup of OxLDL are among the main events that lead to plaque formation.

Clinical presentation in patients with atherosclerosis mostly remains asymptomatic during the early stages of plaque formation; thus, the disease often gets diagnosed only after it has progressed to advanced stages. Once diagnosed, it is not uncommon to see patients having multiple plaques that are in different stages of development and expansion. Currently, once plaques are formed, there are no treatment strategies in place that can regress the developed plaques. Thus, intercepting the early-stage mechanisms of foam cell formation can be helpful in limiting disease progression. Additionally, understanding the signaling mechanisms induced by OxLDL, and the cellular and biochemical pathways affected, is critical to developing effective treatment approaches. Since not all developing plaques will be in the same growth stage, and since plaque formation is a continuous process once initiated, intercepting early-stage mechanisms can prevent the development of new plaques. Some mechanisms that are initiated by OxLDL in macrophages have been previously reported in scientific literature ^6^.

Epigenetic mediators are defined as regulators of gene expression that work through modification to DNA, histones, and other DNA-associated proteins. These epigenetic proteins act as initiators of different biochemical mechanisms in cells. Currently, one of the actively researched topics in heart diseases is investigating the epigenetic roles of methylases, demethylases, and histone deacetylases. In the present work, we investigated the role of TET (Ten Eleven Translocation) and HDAC (Histone Deacetylase) proteins in initiating molecular mechanisms that lead to the formation of atherosclerotic plaques. The findings in this study highlight the early-stage role of TET1 in inducing molecular mechanisms that promote foam cell formation. This study quantitatively measured changes in the expression levels of epigenetic modulators (TETs and HDACs), along with percentage foam cells formed, under TET1 inhibition. These mechanisms presumably are active during the early stages of plaque formation. Furthermore, this study also revealed a possible role of autophagy in foam cell formation, which can also be mediated through epigenetic mechanisms.

## 2 Materials

The following materials, instruments, and software were used: U937 lymphocytes (origin: *Homo sapiens;* ATCC CRL-1593.2); Phorbol 12-myristate 13-acetate (PMA; Sigma-Aldrich P1585), Bobcat339 (Sigma-Aldrich SML2611-5MG), Romidepsin (MedChemExpress HY-15149), RPMI-1640 cell culture medium, Fetal Bovine Serum, 100X antibiotic-antimycotic (ThermoFisher 15240062), DPBS, Takara PrimeScript 1^st^ strand cDNA synthesis kit (Takara catalog no. 6110A), Power SYBR green PCR master mix (Applied Biosystems, catalog no. 4367659), Oil Red O staining kit (ScienCell 0843), Oxidized LDL (NC0955945 Fischer Scientific), RNeasy mini kit (Qiagen, catalog no. 74104), Barrier tips, Eppendorf tubes, 4-well Chamber slides, T25 and T75 flasks (charged plate, perforated lid), 50 ml conical tubes.

Gene-specific primers were designed using NCBI resources and purchased from IDT. Primer sequences for qPCR are as follows:

18Sf: 5’ CCCCTCGATGCTCTTAGCTG 3’

18Sr: 5’ GAACCGCGGTCCTATTCCAT 3’

TET1f: 5’ CAAGTGTTGCTGCTGTCAGG 3’

TET1r: 5’ AATTGGACACCCATGAGAGC 3’

TET2f: 5’ CCAATAGGACATGATCCAGG 3’

TET2r: 5’ TCTGGATGAGCTCTCTCAGG 3’

HDAC2f: 5’ GAGCTGTGAAGTTAAACCGACA 3’

HDAC2r: 5’ ACCGTCATTACACGATCTGTT G 3’

HDAC10f: 5’ ACTGCACTTGGGAAGCTCCTGTA 3’

HDAC10r: 5’ GCCTCTCCGAACAGCCACAT 3’

PyMOL was used to assess protein structure interactions between the epigenetic mediators. The RCSB database was used to obtain protein .pdb files of TET1 (5CG9), TET2 (4NM6), HDAC2 (4LXZ) and HDAC10 (5TD7).

## 3 Methods

### 3.1 Cell culture

#### 3.1a: Revival and culture of U937 cells

The obtained frozen U937 cells were thawed by swirling the vial in a 37°C water bath for 60-90 seconds so that little to no frozen cryopreserving media remained. The thawed cell suspension was added to a 50 ml conical tube that contained complete media (warm, sterile-filtered: 89% RPMI-1640, 10% FBS, 1% 100X antibiotic-antimycotic). The tube was centrifuged at 22°C, 3000 rpm, for 5 minutes. The supernatant was discarded, and the cell pellet was resuspended in 1 ml fresh complete media. The suspension was transferred to a T75 flask containing 9 ml complete media, and placed in an incubator that was maintained at 37°C and 5% CO_2_. After 5 days, cell counts were performed using a hemocytometer and about 0.1 million cells were transferred to a fresh T25 flask containing 5 ml complete media. This step was repeated again after 3 days (once confluence was reached and the color of the culture medium changed). For cytological observations, about 50K cells in 1 ml of culture medium were seeded in each well of a 4-well chamber slide. Once seeded, these cells were immediately differentiated into macrophages with PMA as described in section 3.1c.

#### 3.1b: Isolation of Peripheral Blood Mononuclear Cells (PBMCs) from porcine blood

By accessing the jugular vein, about 20 ml of fresh blood from a healthy pig was collected into an EDTA blood tube. The blood was mixed with an equal volume of sterile PBS. About 20 ml of room-temperature Histopaque 1077 was transferred to a 50 ml conical tube. The blood-PBS mixture was carefully pipetted on top of the Histopaque, making sure that a divide remained between the two substances, and the tube was centrifuged at 400g for 30 minutes, without brakes. After centrifugation, the buffy coat present in the gradient interface was carefully collected and transferred to a 50 ml conical tube containing 35 ml of sterile PBS. The contents in the tube were mixed by gently inverting a few times and then centrifuged at 400g for 5 minutes. The supernatant was aspirated, and the pellet was suspended in 1 ml complete media. The cells were counted using a hemocytometer and about 1 million cells were seeded into each of the T25 flasks containing 5 ml complete media. After overnight incubation at 37°C with 5% CO_2_, the unbound cells were aspirated and the bound cells, which are mostly monocytes and macrophages, were used for Romidepsin treatment (section 3.1d).

#### 3.1c: U937 cell differentiation into macrophages

U937 cells were cultured as suspension cells in T25 flasks containing 5 ml complete media. Their differentiation into macrophages and foam cells was carried out either in T25 flasks or in 4-well chamber slides. To differentiate U937 monocytes into monocyte-derived macrophages (MDMs), 100 ng/ml PMA was added to the culture media and incubated for 48 hours at 5% CO_2_ and 37°C. PMA treatment was repeated every 48 hours for the duration of the cell culture experiment.

#### 3.1d: Treatment with Romidepsin (HDAC2 inhibitor)

Romidepsin has been shown to selectively inhibit HDAC2 at 47 nM ^7^. Since Romidepsin was reported to exert anti-leukemic effects on U937 cells ^8^, primary monocytes/macrophages that were isolated from pig blood were used instead. A total of one million PBMCs isolated via density gradient separation, as described above (section 3.1b), were seeded in T25 flasks containing 5 ml complete media. After overnight incubation at 37°C and 5% CO_2_, the supernatant and unbound suspended cells were aspirated off, and fresh complete media was added. The adherent cells that remained were primarily monocytes and macrophages. These adherent cells were treated for 72 hours with Romidepsin at a final concentration of 47 nM. Total RNA from the control and treated samples was isolated (section 3.2) and RT-qPCR was carried out as described below (section 3.3), to analyze the monocytes/macrophages’ expression of different epigenetic mediators.

#### 3.1e: Treatment with Bobcat339 (TET1 inhibitor)

U937 cells were first differentiated into macrophages (section 3.1c). Bobcat339 was dissolved in DMSO (using the minimum DMSO volume required to dissolve) and added to the culture medium at a final concentration of 33 µM, the concentration at which TET1 is inhibited in culture. Control cells were treated with only DMSO at an equal volume. Cells were incubated for 24 hours at 5% CO_2_ and 37°C. Cells were treated every 24 hours with Bobcat339 for the duration of the cell culture experiment.

#### 3.1f: Inducing foam cell formation

To initiate foam cell formation, the supernatant and unbound cells was aspirated, and fresh complete media containing 80 µg/ml OxLDL was added to the cultured differentiated macrophages (bound cells). PMA and Bobcat339 were also added to the media, depending on their respective treatment times (PMA: every 48 hours; Bobcat339: every 24 hours). The cells were then incubated for an additional 48 hours at 5% CO_2_ and 37°C to allow OxLDL uptake and subsequent foam cell formation. OxLDL treatment was only done after cells had been differentiated and treated with drug inhibitors. Upon completion of the 48 hour incubation with OxLDL, foam cell formation was evaluated under a light microscope before RNA isolation.

### 3.2: RNA isolation

Once cell culture treatments were complete, media was aspirated and the adherent cells were rinsed with DPBS, which was then aspirated. 600µl of Buffer RLT was added evenly across the flask, which was placed on ice for 15 minutes. Using a cell scraper, the lysate was pooled; 6 µl β-mercaptoethanol (10 µl per 1000 µl Buffer RLT) was added to the lysate, and the lysate was aliquoted and stored at -80°C until RNA isolation.

RNA isolation was conducted following the QIAGEN RNeasy Mini Kit protocol for”Purification of Total RNA from Animal Cells Using Spin Technology”. Lysate was thawed on ice and transferred onto a QIAshredder column, then centrifuged on a tabletop centrifuge at 13,000 rpm for 2 minutes. To the collected homogenized flow-through, an equal volume of 70% ethanol was added. After subsequent wash and elution steps according to protocol, 50 µl of RNase-free water was added to the dried column and incubated at room temperature for 5 minutes. The column was then centrifuged at 13,000 rpm for 1 minute, and the flow-through (containing the eluted RNA) was collected. RNA yield was measured using a Nanodrop spectrophotometer, and RNA was stored at -80°C until further use.

### 3.3: Reverse Transcriptase Quantitative Polymerase Chain Reaction (RT-qPCR)

#### 3.3a: First strand synthesis (cDNA synthesis)

cDNA was synthesized following the Takara PrimeScript 1st Strand cDNA Synthesis Standard Protocol. First strand synthesis was a two-step process. First, approximately 5 µg of total RNA was aliquoted into a 100 µl PCR tube, to which 1 µl of Oligo dT primer mix (50 µM) and 1 µl of dNTP mix (contains 10 mM concentration of each of the 4 dNTPs) was added. The total volume was then made up to 10 µl with RNase-free dH_2_O. The mix was incubated at 65°C on a hot plate for 5 minutes to denature the contents, then immediately cooled on ice.

To create the reaction mixture, 4 µl of 5X PrimeScript buffer, 20 units (0.5 μl) of RNase inhibitor, and 200 units (1 μl) of PrimeScript reverse transcriptase were added to the tube, and the reaction mix was made up to a final volume of 20 µl with RNase-free dH_2_O (4.5 μl). After mixing gently, the PCR tube with the reaction contents was incubated at 30°C for 10 minutes, then at 42°C for 60-90 minutes. The tube was then incubated at 70°C for 15 minutes and then immediately cooled on ice. cDNA yield was evaluated using a Nanodrop spectrophotometer, then stored at -20°C until further use. The cDNA synthesized was subsequently used for qPCR amplification of the transcripts of interest expressed.

#### 3.3b: qPCR with SYBR green mix

The reaction mixture containing gene specific primers, cDNA, and SYBR green master mix was made up to a total volume of 12 µl with RNase-free water. This reaction mix underwent qPCR amplification and detection on an Applied Biosystems 7500 Fast Real-Time system. Samples were run in triplicates for each gene and for the control 18S RNA. The thermal cycler conditions were set as: 1) 95°C for 10 minutes for initial denaturation, 2) 95°C for 5 secs followed by 54°C for 1 minute; this step was repeated for 40 times, 3) 95°C for 15 seconds followed by 54°C for 1 minute for dissociation, 4) 95°C for 15 seconds followed by 60°C for 15 seconds for detection. The average Ct values obtained from the triplicate samples were used for measuring the fold change differences, following the ddCt method.

### 3.4: Structural analysis

The interactions between the selected proteins were evaluated on Boston University’s ClusPro server with Piper docking program ^9^. The docking results with balanced coefficients were selected and used for structural visualizations in PyMOL ^10^. Protein-protein interactions were determined by assessing the total number of polar contacts (hydrogen bonds) between the interacting amino acids of the two polypeptide chains from two different proteins. Bond distance was also evaluated; the shorter the bond distance, the stronger the binding between the interacting amino acid residues. The bond distances in Angstroms were determined using PyMOL, and the total number of contacts between the two interacting polypeptide chains of different proteins were counted. Figure 1 below shows the polar contacts and bond distances between the interacting residues of two proteins. The number of hydrogen bonds, and the average bond distance between the two interacting proteins, were shown in graphical format.

**Figure 1:**
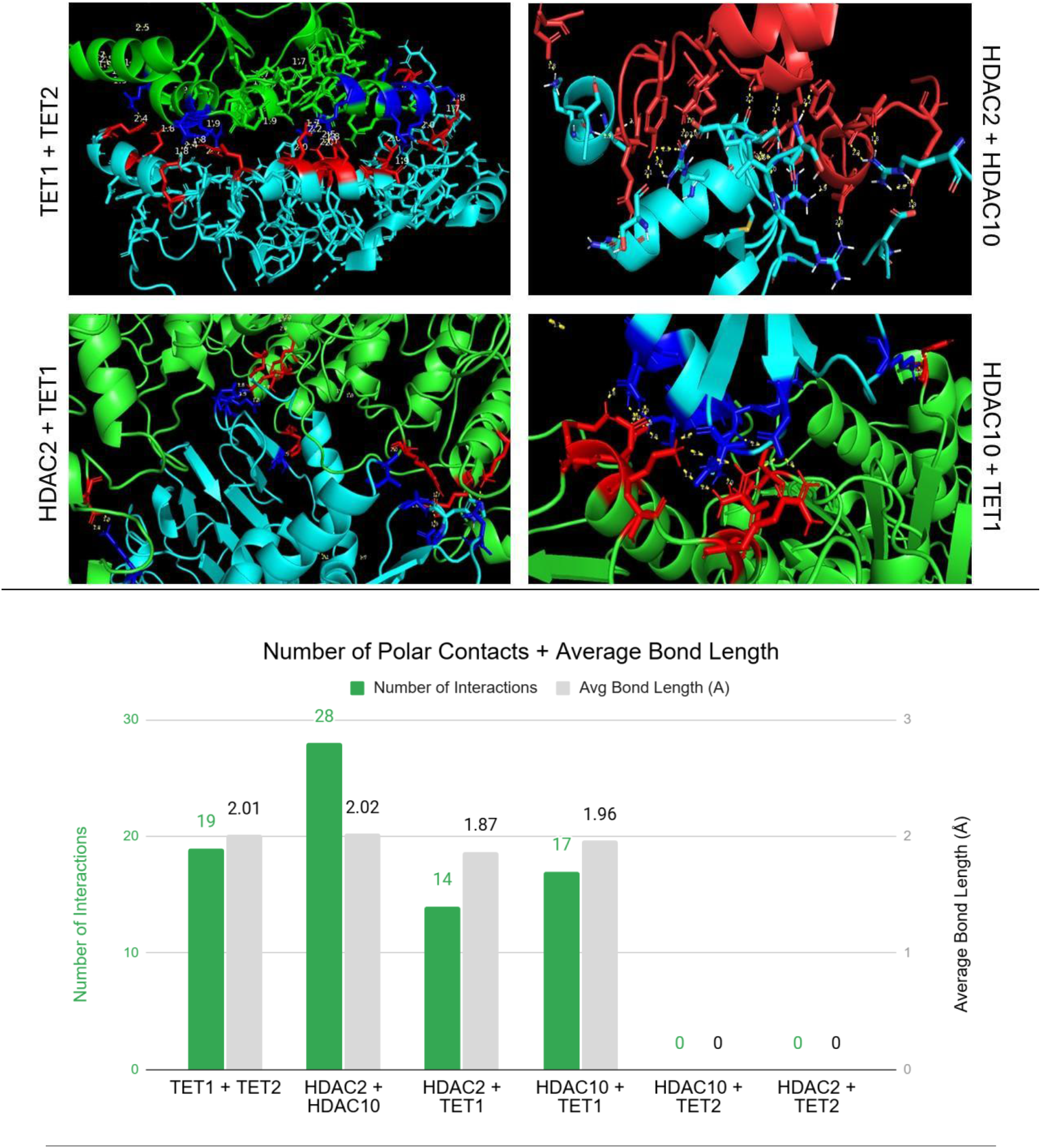
Structural interactions between TET1, TET2, HDAC2, HDAC10. Yellow dotted lines signify polar contacts. Red and blue residues signify interacting residues.

### 3.5: Statistical analysis

Fiji, which is an advanced version of NIH ImageJ software, was used to assess foam cell formation ^11^. Multiple brightfield images of cells in control and different treatment groups were randomly acquired using a Leica DM IL inverted light microscope. The acquired images were then uploaded into Fiji and evaluated by setting the flowing parameters, as follows: 1) Each image was first converted to an 8-bit image and the threshold was adjusted to exclude the background. 2) The images were then converted to Binary, and the scale was matched to the micrometer scale bar. A diameter of 14-micron length, which is equivalent to the diameter of an undifferentiated U937 cell, was considered. For detecting macrophages, the particle size was set to include diameters from 25 microns to infinity. Analysis output was selected to provide Ferrets perimeter and area. Considering that U937 monocytes have a radius (r) of about 7 microns, cells with a perimeter (2πr) of >44 microns and an area (πr^2^) of >154 were considered as representing macrophages/foam cells.

The results obtained from Fiji analysis were saved as a .csv file and statistical analysis was performed in Microsoft Excel. Based on the above cutoff perimeter and area values, the number of macrophages/foam cells formed in control and treatment groups were quantified. The number of foam cells formed in each group were scored, and the mean values with error bars corresponding to standard error of mean (SEM) were represented in bar-graph format. The statistical differences between the groups were assessed using Student’s t-test, and a p-value of less than 0.05 between any two groups was considered as statistically significant.

## 4: Results

### 4.1: Structural interactions between TET and HDAC proteins

To evaluate the structural associations between HDAC and TET proteins, molecular interactions between these two proteins were assessed by docking the corresponding .pdb files using ClusPro.

The number of polar contacts between the two proteins (hydrogen bonding) and the bond distances between them were analyzed. Figure 1 shows the degree of interactions between the two polypeptide chains. The structural interactions between the two proteins were visualized and quantified using PyMOL. No structural interactions were observed between HDAC10 and TET2, nor between HDAC2 and TET2. The bar graph depicts the number of polar contacts and their average bond length in Angstroms; a higher number of interactions alongside a lower average bond length signifies a closer interaction.

### 4.2: Effects of TET1 inhibition on gene expression of epigenetic mediators in macrophages and foam cells

TET1 has been reported to elevate cellular inflammation by upregulating the expression of proinflammatory markers IL1b, IL6, IL8, and TNF-α ^12–15^. Here, the effects of TET1 inhibition in macrophages and foam cells, and TET1’s role in regulating the expression of other epigenetic mediators, were evaluated. Figure 2 shows the expression patterns of different epigenetic regulators that were analyzed using RT-qPCR. In U937 monocyte-derived macrophages (MDMs), TET1 inhibition was seen to decrease the expression of HDAC2. In foam cells, TET1 inhibition not only caused a decrease in the expression of HDAC2 but it also led to decrease in the expression of TET2. ^11^

**Figure 2:**
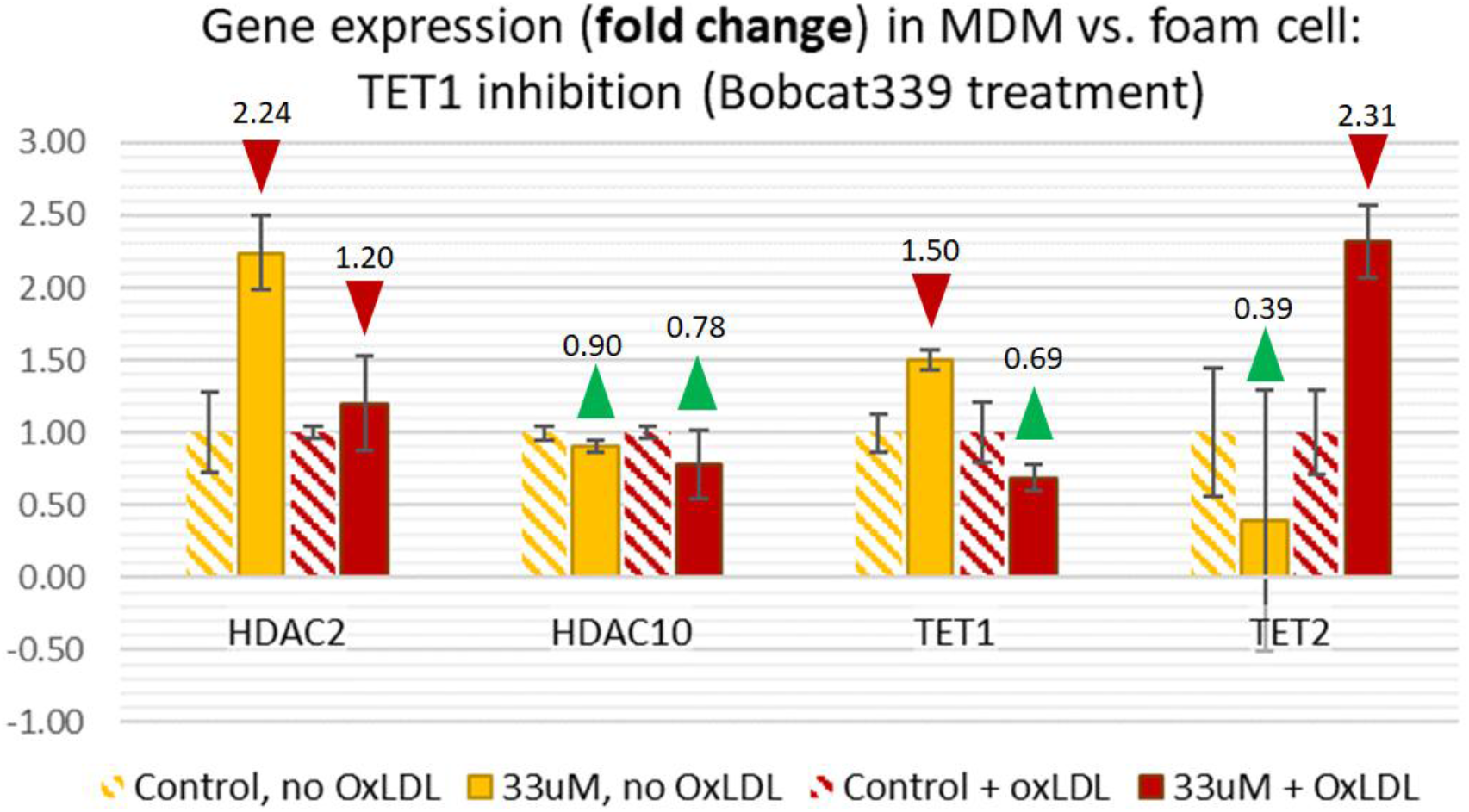
Fold change of target transcripts in PMA-induced monocyte-derived macrophages (MDMs) and OxLDL-treated MDMs (foam cells) in control vs. TET1-inhibited conditions. Here, fold change method was used, which shows the magnitude of change in Ct without information on whether the change is positive or negative. Red/green arrows are included to indicate whether the original dCt change was positive (green; increase in Ct) or negative (red; decrease in Ct).

### 4.3: Effect of HDAC2 inhibition on the expression of epigenetic mediators in porcine macrophages

Romidepsin (HDAC2-specific inhibitor) treatment was reported to confer anti-leukemic effects on U937 cells ^17,18^. Thus, primary monocytes and macrophages were cultured and tested here. Expression of TET and HDAC proteins were analyzed by RT-qPCR. Figure 3 shows that inhibition of HDAC2 leads to a decrease in the expression (measured via fold change, ddCt method) of TET1, TET2 and HDAC10, compared to control.

**Figure 3:**
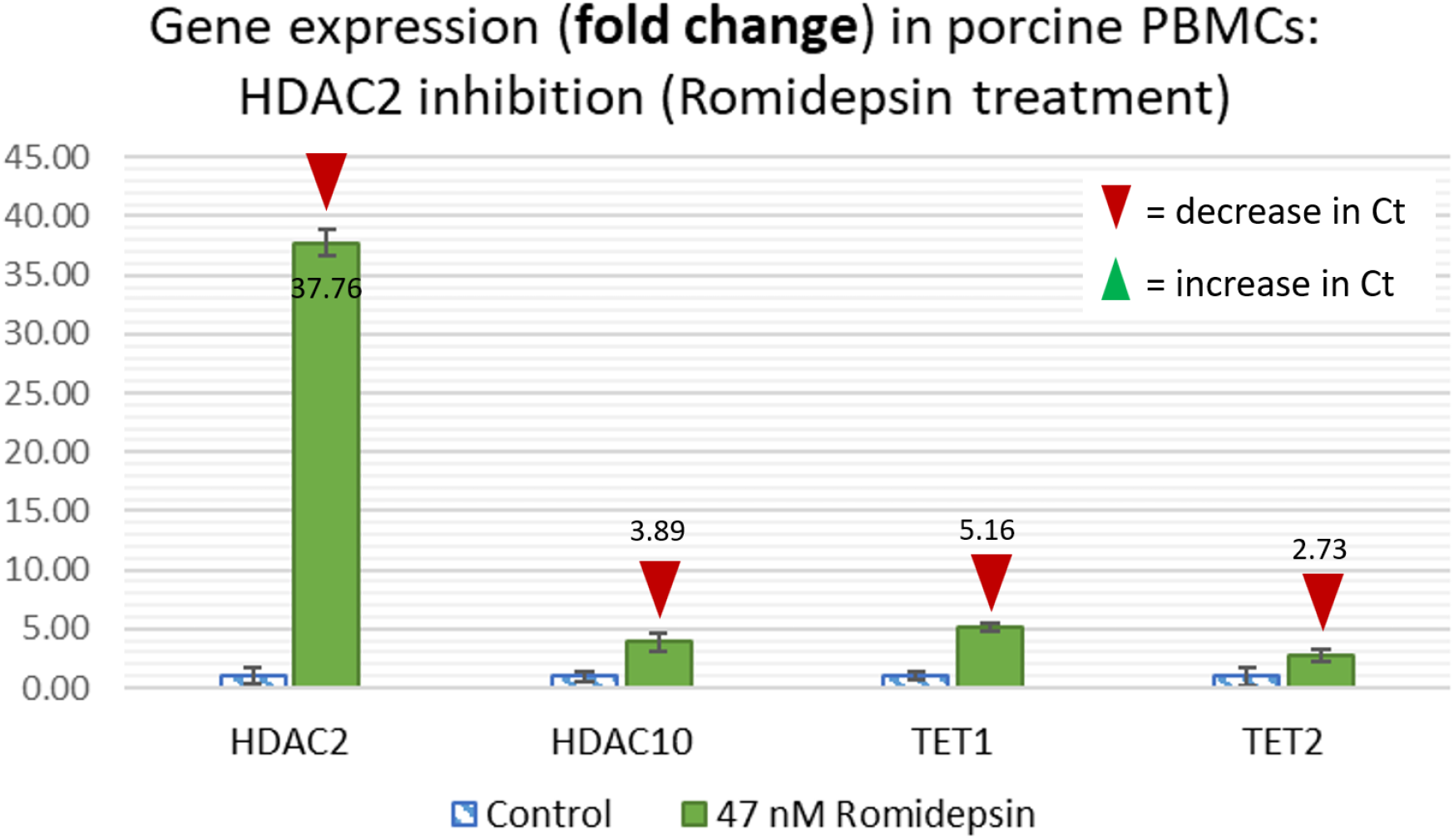
Fold change of target transcripts in porcine peripheral blood macrophages and monocytes (PBMCs) that were untreated (control) vs HDAC2-inhibited (Romidepsin). Here the fold change method was used, which shows the magnitude of change in Ct without information on whether the change is positive or negative. Red/green arrows are included to indicate whether the original dCt change was positive (green; increase in Ct) or negative (red; decrease in Ct).

### 4.4: Effects of TET1 inhibition on foam cell formation

At 33 µM concentration, Bobcat339 was reported to function as a selective inhibitor of TET1 ^16^. During differentiation of THP-1 cells into macrophages with PMA, cellular inflammation was seen elevated by TNF-α. Knock down of TET1 in THP-1 cells was shown to reduce the expression of TNF-α even in presence of the inflammation inducer, LPS ^13^. Furthermore, human monocytes and macrophages were reported to release TNF-α when treated with OxLDL ^19^. Here the effect of TET1 inhibition on OxLDL-induced inflammation was tested. The number of foam cells in different treatment groups were imaged in bright fields and quantified using ImageJ software. Quantitative differences in the percentage of foam cell formation in different treatment groups were graphically represented in Figure 4. Percent foam cell formation was determined by dividing the foam cell count by the total cell count (consisting of both foam cells and macrophages). Cell counts were averaged over 6 image fields.

**Figure 4:**
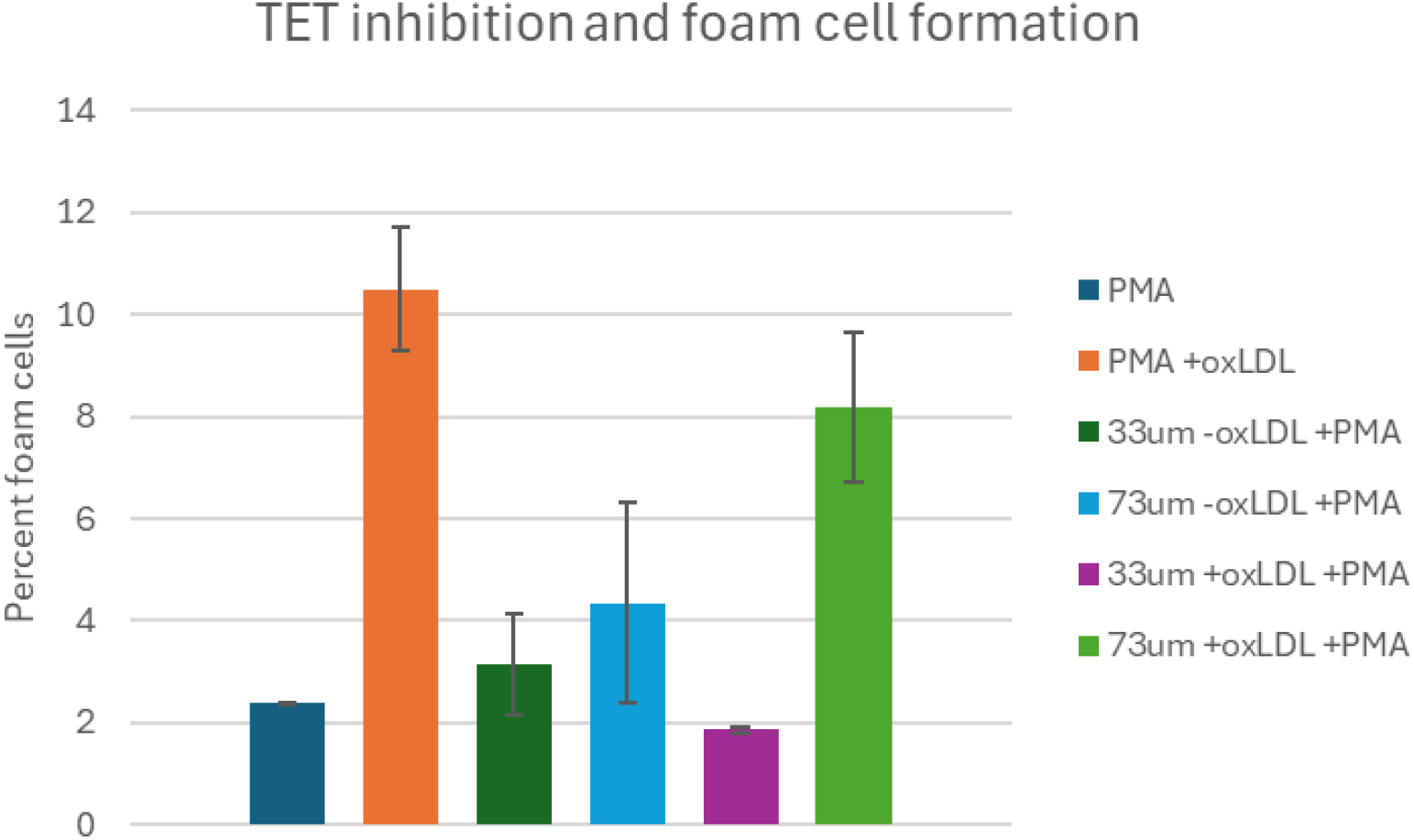
Percent foam cells (of total cells) under TET inhibition conditions. TET1 is inhibited at 33 µM, and TET2 is inhibited at 73 µM. “-oxLDL +PMA” signifies macrophages, and “+oxLDL +PMA” signifies macrophages treated with OxLDL (foam cells).

### 4.5: Role of HDAC10 in foam cells

In macrophages and foam cells that were treated with or without TET1 inhibitor, no significant change in the expression levels of HDAC10 were observed (Figure 2). Also, TET2 does not appear to structurally interact with HDAC10 (Figure 1). It is still not clear whether TET2 has any role in regulating HDAC10 activity during foam cell formation. However, based on our current analysis TET1 appears to interact with other epigenetic mediators and influences their expression. In Figure 5, we illustrate a possible molecular pathway involving TET and HDAC epigenetic mediators during foam cell formation.

## 5: Discussion

Both IL6 and TNF-α not only induces the production of acute-phase reactants, but their induced molecular mechanisms can also cause sustained chronic inflammation in cells. In relevance to atherosclerosis, these proinflammatory agents initiate mechanisms that favor intracellular accumulation of OxLDL, which leads to foam cell formation. Although foam cell formation is speculated to be mediated by multiple mechanisms in several cell types, one proven mechanism involves OxLDL uptake mediated by scavenger receptors, which was demonstrated using THP-1 derived macrophages ^20^. By treating the THP-1 macrophages with FITC (fluorescent dye)-conjugated OxLDL, OxLDL’s role in inducing the production of IL6 and TNF-α was delineated^21^. Different epigenetic mechanisms have been reported to regulate the expression of such inflammation mediators. In THP-1 derived macrophages (another cell line commonly used in studying macrophage functions and atherosclerosis), knockdown of TET1 was found to significantly reduce the expression of TNF-α, which is a key inducer of cellular inflammation ^13^. On the contrary, TET2, which is also one of the isoforms of TET proteins, was identified to associate with HDAC2 and facilitate their binding to the promoter region of IL6, to repress its expression. This was demonstrated in TET2 knockout mice, which had increased IL6 expression levels when challenged with LPS ^22^. In the same article, it was also reported that TET2 affects the expression of IL6 but not TNF-α. Furthermore, during foam cell formation, both IL6 and TNF-α were identified to play critical roles in elevating cellular inflammation. They also initiate different biochemical pathways in macrophages ^20^.

Either TET1 or TET2 deficiency in mice favored the progression of atherosclerosis; however, the molecular functions by which these two proteins promote atherosclerosis were different ^23,24^. Human TET1 and TET2 proteins share less than 65% of sequence similarity with mouse TET1 and TET2. It also appears that TET1 and TET2 may have distinct and yet opposing roles in regulating immune responses in macrophages. As atherosclerosis is primarily an immune disorder, in the present work we examined the role of TET proteins and their correlations with the expression of HDAC proteins in 1) U937 cells derived macrophages, 2) primary porcine peripheral blood macrophages, and 3) induced foam cells (macrophages treated with OxLDL).

The association between TET2 and HDAC2 in repressing IL6 was shown in bone marrow derived macrophages from TET2-deficient mice ^22^. However, this association may not be on the structural level (not related to the protein-protein interactions between them). A recent study has shown that HDAC2 does not bind to the promoter regions of proinflammatory genes, but knockdown of HDAC2 may cause a decrease in the production of inflammatory markers ^25^. We believe that since both TET2 and HDAC2 are transcriptional regulators, these two proteins may be part of a larger transcriptional machinery involved, and their mere association requires other proteins in the transcriptional complex to mediate. To determine whether a structural interaction between HDAC2 and TET2 is feasible, we conducted structural analysis via PyMOL. Our analysis did not identify any hydrogen bonding to exist between HDAC2 and TET2. Subsequently, we extended our analysis and assessed protein structural interactions between TET1, TET2, HDAC2 and HDAC10. In Figure 1, along with the structures of the interacting proteins, we show the number of hydrogen bonds between the interacting residues and their average bond distances. Our analysis shows that TET1 seems to structurally interact with TET2, HDAC2 and HDAC10, but TET2 interacts only with TET1—not with HDAC2 and HDAC10. In addition to this, structural interactions between HDAC2 and HDAC10 were also observed.

Since TET1 was found to interact with the other three epigenetic regulators (TET2, HDAC2, HDAC10) that are described above, we examined the importance of TET1 in U937-derived macrophages and foam cells. We evaluated the variations in the transcript levels of TET1, TET2, HDAC2, and HDAC10 by treating cells with or without 33 µM Bobcat339, a specific inhibitor of TET1 (Figure 2). As anticipated, treatment with the TET1 inhibitor greatly reduced the transcript levels of TET1 in macrophages. However, in OxLDL-induced foam cells that were treated with or without TET1 inhibitor, no significant differences in TET1 transcript levels were observed. This suggests that OxLDL-mediated signaling may initiate direct or indirect mechanisms in foam cells to induce inflammatory responses that are independent of TET1. Our results also show that upon inhibition of TET1 in macrophages, the expression levels of HDAC2 drastically declined, compared to untreated macrophages. The transcript levels of HDAC2 were also decreased significantly in foam cells when TET1 was inhibited. Interestingly, the transcript levels of HDAC2 in OxLDL-induced foam cells were found to be similar to untreated control macrophages. This observation suggests that OxLDL-mediated mechanisms are likely to be functionally independent of HDAC2 in foam cells, and these molecular mechanisms are not active in macrophages. With regard to TET2 expression, no significant differences in its expression levels were observed in macrophages that were treated with or without TET1 inhibitor, nor in foam cells that were not treated with TET1 inhibitor. Noticeably, in foam cells that were treated with TET1 inhibitor, a drastic decrease in the transcript levels of TET2 was observed, suggesting that OxLDL-induced mechanisms may prevent TET2 expression in foam cells, in absence of TET1. These findings also indicate an inter-regulated expression between TET1 and TET2 that occurs only in foam cells. In these results, since TET1 inhibition in foam cells leads to concomitant decrease in the expression levels of both HDAC2 and TET2, our results partly support the discussed published report ^22^ that describes the association between TET2 and HDAC2, but not the structural interactions between them.

Seeing that HDAC2 expression was affected only in macrophages and foam cells that were treated with TET1 inhibitor, we questioned whether HDAC2 inhibition would have an effect on the expression of TET1 and/or on other mediators with which TET1 potentially interacts during foam cell formation. To address this, we tested the effects of Romidepsin (FK228, a specific inhibitor of HDAC2) in porcine macrophages. U937-derived macrophages could not be used for Romidepsin treatment as this drug was reported to cause cell cycle arrest at G1 and G2/M phases in U937 cells by inhibiting HDAC2 ^8^. As seen in Figure 3, Romidepsin treatment to porcine primary macrophages affected the expression of TET1, TET2, and HDAC10, in addition to HDAC2, suggesting that both HDAC2 and TET1 may have a critical role in foam cell formation.

One of the extensively studied mouse models of atherosclerosis is the ApoE knock-out model. In this mouse model, TET2 was reported to inhibit progression of atherosclerosis by upregulating autophagy in vascular endothelial cells ^26^. Supporting this report, in THP-1 derived macrophages that were treated with OxLDL, autophagy markers were seen to be inhibited in absence of TET2, indicating a prominent role of TET2 in preventing atherosclerosis ^27,28^. In the present study, similar results were seen in OxLDL-induced foam cells that were treated with TET1 inhibitor, where a drastic decrease in the expression levels of TET2 was observed (Figure 2). Decreased expression of TET2 in absence of TET1 also correlated with a reduced number of foam cells in this group. Figure 4 shows percent foam cells in different treatment groups. Our results suggest that TET1 inhibition leads to a decreased percentage of foam cells; this correlates with the decreased expression of HDAC2 and TET2, further supporting a previous report that described an association between HDAC2 and TET2.

A discern role of HDAC10 in cellular autophagy was reported previously ^29,30^. A recent report, describing the development of two HDAC10-specific inhibitors, demonstrated that specific inhibition of HDAC10 would promote cellular autophagy^31^. In later stages of atherosclerosis when plaques are fully formed, foam cell formation in VSMCs was observed to be elevated, and this correlated with inhibition of cellular autophagy ^2^. Inhibition of autophagy could lead to destabilization of the atherosclerotic plaques that are in advanced stages of growth. Thus, inducing autophagy using pharmacological agents has been investigated, as a potential help in stabilizing the vulnerable plaques ^32,33^. Since TET2 confers inhibitory effects on foam cell formation, we are currently investigating whether it has an indirect role in initiating HDAC10-mediated autophagy. The direct role of TET2 in regulating HDAC10 appears to be not feasible, as we show here (Figure 1) that these two proteins do not interact structurally. In agreement with the above reports and our observations, we speculate that the indirect effects of TET2, if any, in HDAC10-mediated autophagy may not be part of the early-stage mechanisms occurring in foam cells. It is to be noted that in lung pathologies, HDAC10 seems to promote inflammation in macrophages and its knockdown confers protection from acute lung injury ^34–36^. Based on our analysis, a unique role of HDAC10 is yet to be deciphered, which can project its role in promoting inflammation, preventing autophagy and inducing foam cell formation.

## 6: Conclusions and unique findings

In conclusion, studies in this manuscript indicate that: **1)** TET2 structurally interacts only with TET1, but not with HDAC2 and HDAC10. However, TET1 interacts with TET2, HDAC2 and HDAC10. **2)** In U937-derived macrophages and in OxLDL-induced foam cells, inhibition of TET1 correlates with decreased expression of HDAC2. **3)** OxLDL appears to have no role in inducing the expression of HDAC2 in foam cells. **4)** While TET1 inhibition has no effect on TET2 in macrophages, in OxLDL-induced foam cells, TET1 inhibition greatly affects the expression of TET2. **5)** TET2 may have a late secondary role in promoting HDAC10-mediated autophagy in foam cells. To our knowledge, these are the first reports that show the role of TET1 in promoting foam cell formation, and its associated molecular mechanisms with HDAC2, TET2 and HDAC10.

## 7: Acknowledgements

We thank Benjamin Nelson for providing training and guidance with routine cell culture methods, instrument usage, and overall workflow in the laboratory. Funding for this project was supported by the MizzouForward Undergraduate Research Training Grant (2024), Peggy & Andrew Cherng Summer Scholarship (2023), and Momentum Grant (2021) to AB, and departmental funds to JG.

## 8: Abbreviations

TET1: Ten Eleven Translocation 1
TET2: Ten Eleven Translocation 2
HDAC2: Histone Deacetylase 2
HDAC10: Histone Deacetylase 10
OxLDL: Oxidized Low Density Lipoproteins
PMA: Phorbol 13-Myristate 12-Acetate
FBS: Fetal Bovine Serum
PBMCs: Peripheral Blood Mononuclear Cells
NBF: Neutral Buffered Formalin

